# Characterizing insect communities within thin-soil environments

**DOI:** 10.1101/2022.10.01.510429

**Authors:** Katherine McNamara Manning, Kayla I. Perry, Christie A. Bahlai

**Affiliations:** Department of Biological Sciences, Kent State University, Kent, Ohio, USA; Department of Entomology, Ohio State University, Wooster, Ohio, USA

## Abstract

Natural thin-soil environments are those which have little to no soil accumulation atop hard substrates. Many of these natural thin-soil environments, such as alvars, rocky lakeshores or glades, cliffs and cliff bluffs, and barrens, are found in the Great Lakes Region of North America. Due to their ubiquity and ecosystem services they provide, characterizing insects in sensitive environments such as these is important. This study monitored insects in nine thin-soil sites, within three regions, on a 630 km latitudinal gradient in the Southeastern Great Lakes Region of North America from June - August 2019. Over 22,000 insect specimens collected were identified to order or family, and bee specimens were identified to genus or species. We found that overall insect community composition and biodiversity characteristics were similar between the three regions examined. However, the central region had higher taxonomic richness than the southern region. Although unique bee taxa were observed in each region, diversity metrics and community composition of bees were similar among sites. This study provides taxonomic information about the insect, particularly bees, and plant communities in thin-soil environments in this region, which could support conservation and management efforts.

## Introduction

Ecological communities are shaped by the physical attributes of their environments. In the Great Lakes basin, numerous globally rare habitats occur, including those characterized by their paucity of surface soil. The ecological class ‘Primary’ is characterized by having little to no soil accumulation on top of bedrock, cobble, or exposed mineral soil. There are many natural thin-soil community types, such as alvars, rocky (cobble or bedrock) lakeshores or glades, cliffs and cliff bluffs, and barrens (Cohen et al. 2015). These habitats may experience intense wind and solar radiation, as well as varying precipitation conditions, from heavy rain to drought, due to the bedrock being at or near the surface (Stephenson and Herendeen 1986, Brunton 1988, Albert 2006). Together, these factors limit the primary producers that can survive there (Lundholm 2006), creating naturally open landscapes typically with low growing vegetation (Reschke et al. 1999). These unique and sensitive habitats are often home to rare plant and insect species adapted to these unusual environments, making them areas of research interest and conservation concern (Comer et al. 1997, Reschke et al. 1999, Albert 2006, Neufeld et al. 2018, McMullin 2019).

In the Great Lakes Region of North America, research in these rare thin-soil environments, mostly alvars, has generally focused on identifying the taxa that occur in them or are unique to them. Some of the characteristic rare vascular plants on Great Lakes alvars include: *Carex juniperorum* W. J. Crins (Juniper Sedge), *Cirsium hillii* (Canby) (Hill’s Thistle), *Cypripedium arietinum* W. T. Aiton (Ram’s Head Lady Slipper), and *Solidago houghtonii* A. Gray (Houghton’s Goldenrod), as well as *Iris lacustris* Nutt. (Dwarf Lake Iris) and *Tetraneuris herbacea* Greene (Lakeside Daisy), both of which are endemic to the Great Lakes. Insect taxa that have been known to show particular affinity for alvars include: carabid beetles, cicadellids, lepidopterans, orthopteroids, and symphytes (Reschke et al. 1999). In alvar surveys in southern Ontario, three species of ground beetles (Carabidae) were dominant, but rarely collected in other non-alvar parts of the province: *Agonum nutans* (Say), *Chlaenius purpuricolliis* Randall, and *Pterostichus novus* Straneo (Bouchard et al. 1998, 2005).

Insects play a variety of critical roles in ecosystem function and service, one of which is pollination (Losey and Vaughan 2006, Noriega et al. 2018). Plant-pollinator relationships are important for biodiversity in natural and managed ecosystems (Ollerton 2017, Wei et al. 2021). Among insects, certain flies, wasps, beetles, butterflies, and moths perform pollination services, however, bees are considered to be the most efficient insect pollinator (Potts et al. 2016). Both wild and managed bee species are vital to at least 35% of crop production (Klein et al. 2007), which is crucial for human food security (Ollerton 2017). Bees are a diverse group of insects, with over 22,000 species worldwide. In North America, north of Mexico, there are over 4,000 bee species (Wilson and Messinger Carril 2016). However, in recent years, several high-profile studies have reported mass declines of insect taxa around the globe (Fox 2013, Hallmann et al. 2017, Lister and Garcia 2018, Loboda et al. 2018, Seibold et al. 2019, Sánchez-Bayo and Wyckhuys 2019, Wepprich et al. 2019, van Klink et al. 2020, Zattara and Aizen 2020). Bees are not immune to this global loss of biodiversity, with reports of declines in managed (vanEngelsdorp et al. 2008, Ellis et al. 2010, Steinhauer et al. 2014) and wild bee species (Colla and Packer 2008, Grixti et al. 2009, Cameron et al. 2011, Scheper et al. 2014, Koh et al. 2016, Arbetman et al. 2017, Graham et al. 2021). Because of their important roles as pollinators, characterizing bee communities in different environments, such as thin-soil environments, is valuable for management and conservation decisions.

Surveys of insects in Great Lakes thin-soil environments have largely focused on smaller geographical regions, such as the intense sampling of alvars of Ontario, especially the Saugeen (Bruce) Peninsula, and in Michigan near the northern shorelines of Lakes Michigan and Huron (Albert et al. 1994, 1995, Reschke et al. 1999, Bouchard et al. 2005, Albert 2006, Cohen et al. 2015). Few surveys have occurred over the north-south expanses of this ecoregion, and none have explicitly examined the communities of insects across these thin-soil habitats spanning the region. Additionally, to our knowledge bee communities in these thin-soil environments have not been characterized before. To address these knowledge gaps, we conducted a study of the community composition of insects in thin-soil environments in three regions on a latitudinal gradient across the Southeastern Great Lakes Region of North America. In this study, we examined biodiversity metrics of insect communities, including bees as our focal taxa, between northern, central, and southern regions of this area. We predicted that even if habitats and plant communities differed, insect communities would be similar across these three regions because the community will be shaped by the abiotic traits of the landscape.

## Methods

### Site descriptions

This study was conducted in thin-soil environments along a latitudinal gradient spanning 630 km in eastern North America (Figure 2). Along this gradient, three sites were selected on the northern Bruce Peninsula in Ontario, Canada, three sites in Cuyahoga County, Northeast Ohio, USA, and three sites in Hocking County, Southeast Ohio, USA (Figure 1). All sites had thin soils (about 15 cm or less) and typically lacked direct canopy cover, leaving them relatively open and exposed to solar radiation, winds, and ranging precipitation conditions. Permits to collect insects and plants were obtained for all sites in accordance with landholder policy.

**Figure 1:**
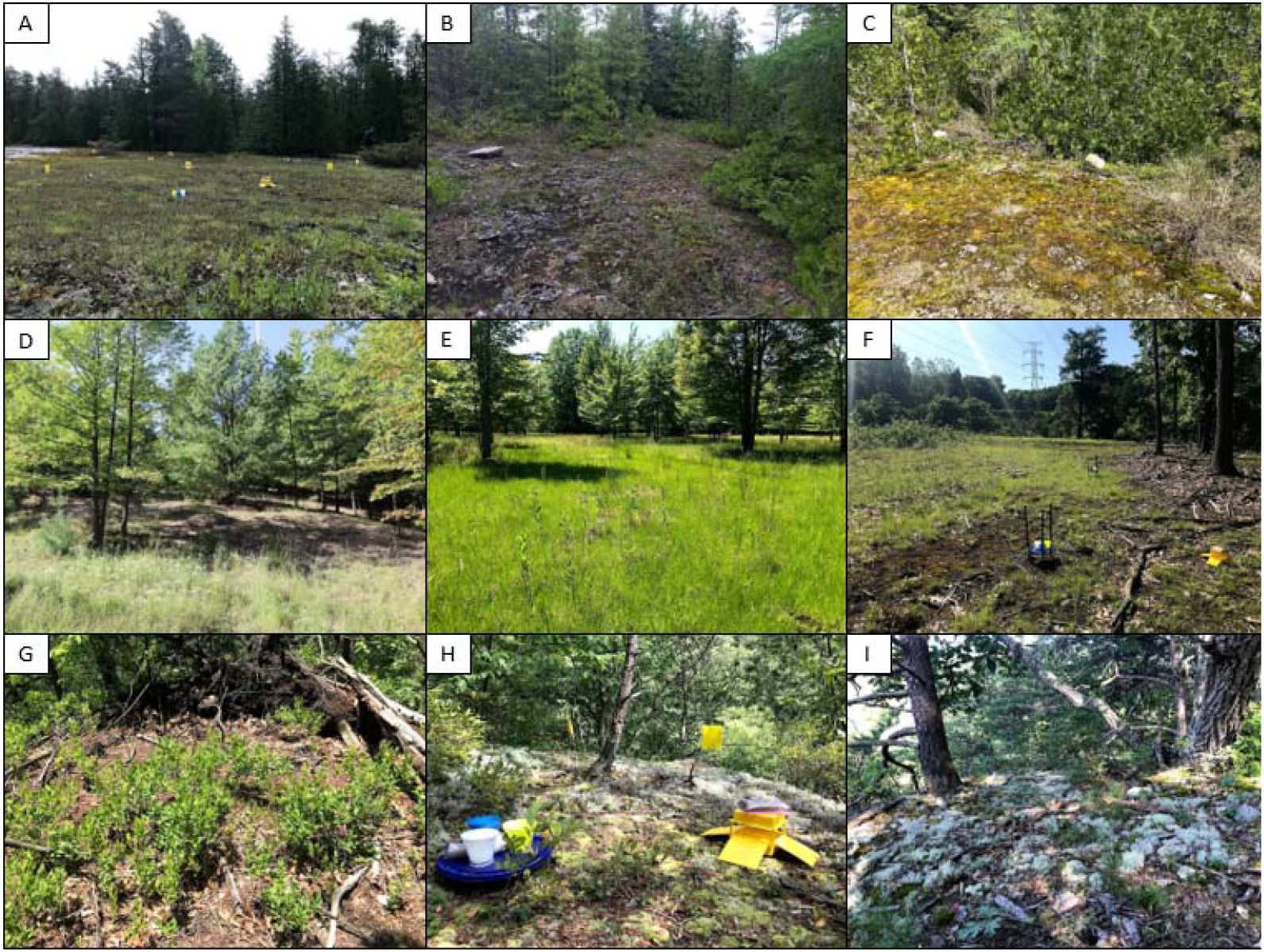
Sampling sites in each region. North: (A) Davis alvar, (B) Beaton alvar, (C) Cape hurd alvar, Central: (D) Slate shale hill, (E) Dusty goldenrod meadow, (F) Bedford barren, and South: (G) Snyder, (H) The “W” – ladder, (I) The “W” – picnic rock.

**Figure 2:**
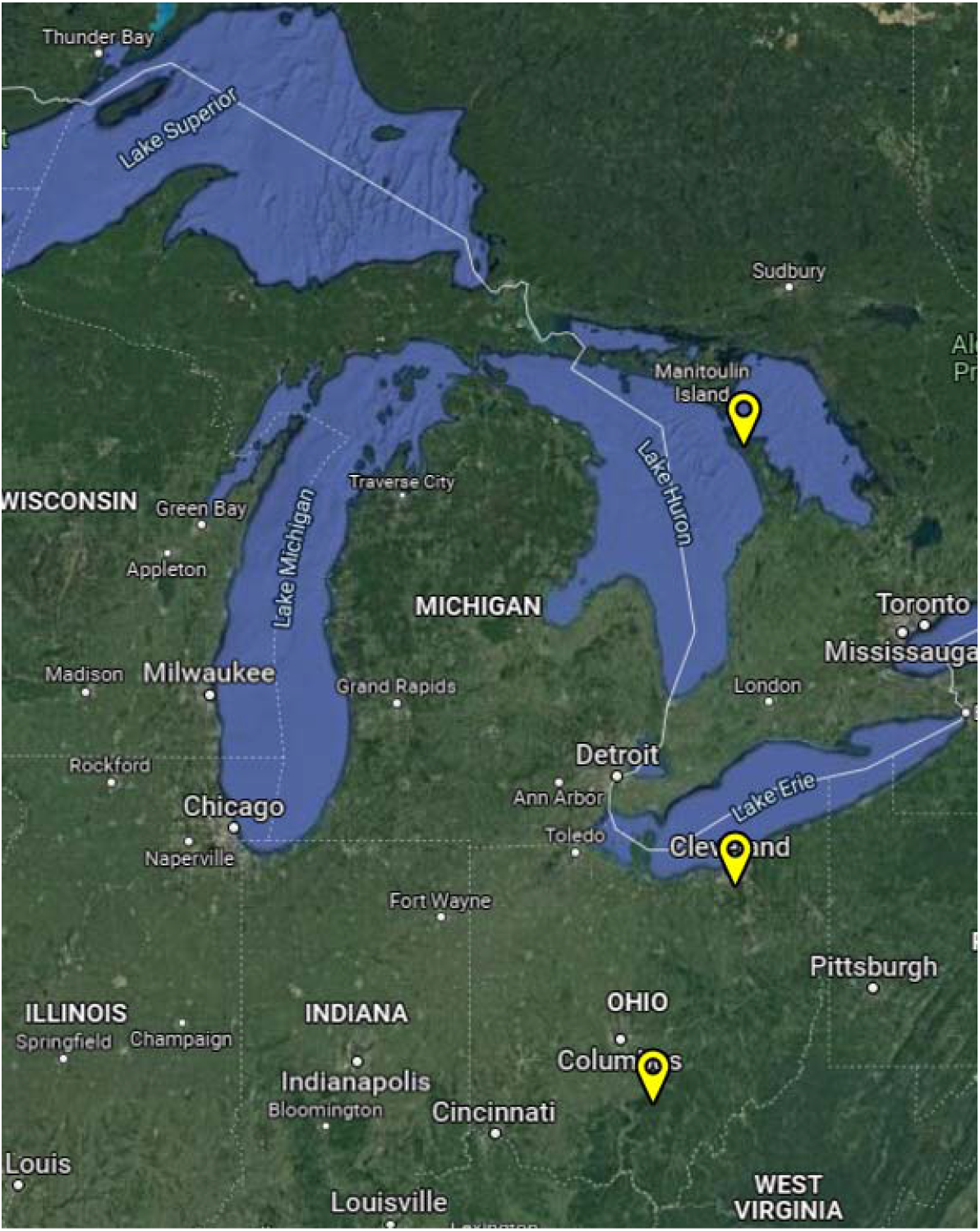
General location of sampling sites in each region. Figure adapted from Google Maps.

Northern sites were located on the Northern Bruce Peninsula, Ontario, Canada along the Niagara escarpment and owned by The Nature Conservancy of Canada. All sites were characterized as alvars, with little to no vegetation on limestone bedrock. The Davis alvar is further defined as a dry lichen-moss open pavement alvar. It had the most open and flat landscape, with sparse, low-growing vegetation. The Beaton alvar was a creeping juniper and shrubby cinquefoil dwarf shrub alvar, with a slightly rocky landscape. The Cape Hurd Alvar was similar to Beaton in vegetation and landscape, but closer to the lakeshore making it subject to frequent low floods. All sites occurred in a rural landscape, primarily consisting of undeveloped land, and protected areas, but the area had previously been subjected to heavy logging pressure.

Central sites were located in Cuyahoga County in Northeast Ohio, USA, specifically in the municipalities of Walton Hills, Parma, and Highland Heights. All sites were embedded in a greater landscape of mixed use, urban, industrial, and semi-urban land use histories. In Walton Hills, we sampled a thin-soil mossy barren near a hiking trail, between a meadow and forest, near a cliff edge over a creek in the Cleveland Metroparks Bedford Reservation. Cleveland Metroparks also owned the site in Parma, which was a roadside hill with slate shale soil, with sparse vegetation and trees. The Highland Heights Dusty Goldenrod Preserve is owned by the West Creek Conservancy, Cuyahoga Soil and Water Conservation District, Friends of Euclid Creek, and Euclid Creek Watershed Council. The portion of the preserve we sampled was an open wet meadow in a forest, with tall vegetation, sparse trees, and is the only known home in Ohio to its namesake, the rare and endangered Dusty Goldenrod (*Solidago puberula* Nutt.). This site had thin soils due to a history of topsoil harvesting.

Southern sites were all located within the 520 hectare Crane Hollow Preserve in Rockbridge, Southeast Ohio, USA, part of the unglaciated Allegheny Plateau. These sites occurred in a highly rural private landscape with sparse human activity and development history. These sites were located on cliff edges on top of deep ravines with a Black Hand sandstone bedrock. The “W” ladder and picnic rocks sites were both named for spots along the “W” hollow and Snyder was along Snyder hollow. All sites were open, xeric, thin-soil areas within forest, dominated by mosses, lichens, and small shrubs. The two “W” sites were heavily covered in Reindeer Lichen (*Cladonia* sp.) and American Wintergreen (*Gaultheria procumbens* L.).

### Field and laboratory methods

The thin soils in these habitats constrained the methods used to monitor insect communities, preventing typical pitfall sampling for ground-dwelling insects from being deployed. From a conservation standpoint, disturbing natural thin soil environments is undesirable, and from a practical standpoint, some of these sites were bedrock with almost no soil. Due to these constraints, all sampling had to be done above the surface level. Insects were surveyed using three types of passive sampling traps: yellow sticky cards (Pherocon, Zoecon, Palo Alto, CA, USA), bee bowls (also known as pan traps, inspired by Leong & Thorp, 1999), and ramp traps (ChemTica Internacional S.A., Santo Domingo, Costa Rica) (Figure 3), evenly spaced at each site for 48 hours once per month in June, July, and August 2019. The spacing and number of traps was dependent on the area available for sampling at the site (Table S1). Each bee bowl was an array of three different colored bowls spray painted fluorescent yellow, blue, or white (Krylon Industrial, Cleveland, OH, USA). Sticky cards and bee bowls were placed at the height of the plant community to collect flying insects, and ramp traps were placed on the surface of the ground to collect ground-dwelling insects. Bowls and ramps were filled with soapy water (Dawn Original Liquid Dish Soap, Procter & Gamble, Cincinnati, OH, USA) and had quart zipper-top bags filled with sand placed on top to minimize wind disturbance. Upon collection, bowls and ramps were strained in the field and samples were placed in a gallon zipper-top bag with 70% ethanol. Sticky cards were placed directly into gallon zipper-top bags.

**Figure 3:**
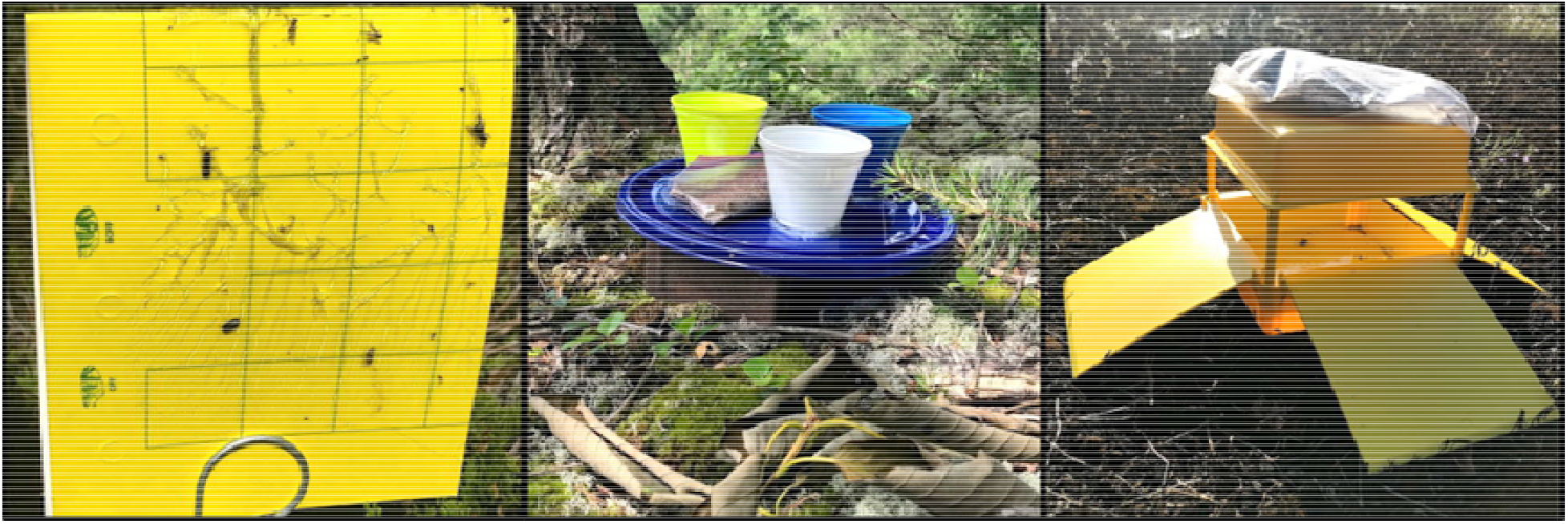
Sticky cards, bee bowls, and ramp traps (left to right) deployed at various field sites in the Southeastern Great Lakes Region in 2019.

Vegetation was sampled at the sites in July-August 2019 by placing a meter-squared quadrat in a well-vegetated location within the site and identifying the vascular plants to species or genus (det. John Gerrath, Joe Moosburgger, and Glenn Vande Water). The trees that directly surrounded the sites, as well as prominent plant species that were not inside the quadrat were also recorded (Table S2).

In the lab, the liquid samples were strained, and specimens were identified and placed in vials with 70% ethanol for storage. The sticky cards were frozen, and specimens were identified while remaining in the bag, afterwards they remained stored in the freezer. Specimens were identified to order, suborder, superfamily, group (“wingless parasitoid wasps”), or family (Table 1). We identified bees (Hymenoptera: Apoidea: Anthophila) to the highest precision possible (genus or species) using expert identification and various interactive Discover Life keys (det. MaLisa Spring, http://www.discoverlife.org).

**Table 1:** Insect abundances, total (T) and standardized (S), by region. Standardized abundances were calculated by dividing total abundances for each taxon by the total number of traps used in that region.

| Order |  | North |  | Central |  | South |  |
| --- | --- | --- | --- | --- | --- | --- | --- |
|  |  | T | S | T | S | T | S |
| Blattodea |  | 0 | 0 | 1 | 0.01 | 0 | 0 |
| Diptera |  | 2510 | 34.86 | 5334 | 74.08 | 1142 | 15.86 |
|  | Syrphidae | 163 | 2.26 | 316 | 4.39 | 21 | 0.29 |
| Ephemeroptera |  | 5 | 0.07 | 0 | 0 | 0 | 0 |
| Hymenoptera |  | 6280 | 87.22 | 687 | 9.54 | 591 | 8.21 |
|  | Anthophila | 15 | 0.21 | 99 | 1.38 | 54 | 0.75 |
|  | Chalcidoidea | 6189 | 85.96 | 433 | 6.01 | 179 | 2.49 |
|  | Chrysidae | 0 | 0 | 0 | 0 | 1 | 0.01 |
|  | Ichneumonoidea | 41 | 0.57 | 104 | 1.44 | 76 | 1.06 |
|  | Dryinidae | 0 | 0 | 1 | 0.01 | 0 | 0.00 |
|  | Formicidae | 28 | 0.39 | 35 | 0.49 | 272 | 3.78 |
|  | Pompilidae | 2 | 0.03 | 0 | 0 | 0 | 0 |
|  | Unknown wasp | 2 | 0.03 | 15 | 0.21 | 9 | 0.13 |
|  | Wingless parasitoid | 3 | 0.04 | 0 | 0 | 0 | 0 |
| Hemiptera |  | 274 | 3.81 | 4681 | 65.01 | 127 | 1.76 |
|  | Aleyrodidae | 2 | 0.03 | 133 | 1.85 | 11 | 0.15 |
|  | Aphididae | 148 | 2.06 | 4317 | 59.96 | 13 | 0.18 |
|  | Cercopidae | 0 | 0 | 7 | 0.10 | 1 | 0.01 |
|  | Cicadellidae | 110 | 1.53 | 163 | 2.26 | 70 | 0.97 |
|  | Fulgoroidea | 0 | 0 | 3 | 0.04 | 0 | 0 |
|  | Membracidae | 0 | 0 | 3 | 0.04 | 11 | 0.15 |
|  | Miridae | 1 | 0.01 | 1 | 0.01 | 0 | 0 |
|  | Orius | 0 | 0 | 1 | 0.01 | 0 | 0 |
|  | Psyllidae | 7 | 0.10 | 24 | 0.33 | 2 | 0.03 |
|  | Tingidae | 0 | 0 | 1 | 0.01 | 0 | 0 |
|  | Unknown Hemipteran | 6 | 0.08 | 28 | 0.39 | 19 | 0.26 |
| Coleoptera |  | 235 | 3.26 | 404 | 5.61 | 188 | 2.61 |
|  | Buprestidae | 0 | 0 | 0 | 0 | 2 | 0.03 |
|  | Cantharidae | 5 | 0.07 | 18 | 0.25 | 1 | 0.01 |
|  | Carabidae | 1 | 0.01 | 1 | 0.01 | 0 | 0 |
|  | Chrysomelidae | 6 | 0.08 | 48 | 0.67 | 2 | 0.03 |
|  | Coccinellidae | 0 | 0 | 1 | 0.01 | 1 | 0.01 |
|  | Curculionidae | 2 | 0.03 | 17 | 0.24 | 4 | 0.06 |
|  | Elateridae | 2 | 0.03 | 1 | 0.01 | 0 | 0 |
|  | Lampyridae | 25 | 0.35 | 1 | 0.01 | 1 | 0.01 |
|  | Meloidae | 1 | 0.01 | 0 | 0 | 1 | 0.01 |
|  | Mordellidae | 8 | 0.11 | 25 | 0.35 | 14 | 0.19 |
|  | Scarabaeidae | 1 | 0.01 | 3 | 0.04 | 5 | 0.07 |
|  | Staphylinidae | 20 | 0.28 | 4 | 0.06 | 32 | 0.44 |
|  | Unknown Coleoptera | 21 | 0.29 | 36 | 0.50 | 18 | 0.25 |
| Lepidoptera |  | 38 | 0.53 | 17 | 0.24 | 1 | 0.01 |
| Lepismatidae |  | 0 | 0 | 0 | 0 | 1 | 0.01 |
| Mecoptera |  | 2 | 0.03 | 0 | 0 | 0 | 0 |
| Neuroptera |  | 0 | 0 | 5 | 0.07 | 0 | 0 |
| Orthoptera |  | 13 | 0.18 | 18 | 0.25 | 12 | 0.17 |
| Psocodea |  | 0 | 0 | 2 | 0.03 | 12 | 0.17 |
| Symphyta |  | 2 | 0.03 | 0 | 0 | 0 | 0 |
| Thysanoptera |  | 79 | 1.10 | 207 | 2.88 | 80 | 1.11 |
| Trichoptera |  | 9 | 0.13 | 0 | 0 | 1 | 0.01 |

### Statistical methods

All statistical analyses were completed using R 4.1.3 (R Core Team, 2022). Model fits were evaluated for statistical assumptions of normality of residuals and homogeneity of variance. Taxonomic richness (number of taxa per sample) and Shannon diversity index (Hill 1973) were calculated using the *vegan 2.5-7* package (Oksanen et al. 2019).

Linear mixed effects models were developed using the *lme4* (Bates et al. 2015) and *lmerTest* (Kuznetsova et al. 2017) packages to examine differences in insects among regions. The response variables examined were taxa richness and Shannon diversity. Each model included region (north, central, south), sampling date (date of trap collection), and trap type (yellow sticky card, bee bowl, ramp trap) as categorical fixed effects and trap number nested within the site as a random effect: *Response variable ~ region+Date+Trap+(1|Site:Replicate)*. The function ‘Anova’ from the car package (Fox and Weisberg 2019) was used to examine the effect of region in each model. Tukey pairwise comparisons were performed using the *emmeans 1.7.4-1* package (Lenth 2021) for all models to compare between regions (north, central, and south).

To visualize the insect communities collected in each region we used non-metric multidimensional scaling (NMDS, with Jaccard distance), computed using the *vegan 2.5-7* package. For this analysis we used presence-absence data pooled by site for each sampling date. Permutational multivariate analysis of variance (PERMANOVA), analysis of multivariate homogeneity of group dispersions (BETADISPER), and pairwise multilevel comparison using the adonis function were performed following the NMDS analysis to assess compositional dissimilarity between regions. NMDS, PERMANOVA, and BETADISPER were computed using functions in the *vegan 2.5-7* package. Pairwise adonis (pairwise PERMANOVA) was performed using the *pairwiseAdonis* package (Martinez Arbizu 2020).

Accumulation curves of bees for each region were created using the *BiodiversityR* package (Kindt and Coe 2005). To estimate sampling efficiency for each trap type, we used nonparametric Jackknife order 1 estimator to compare observed and estimated richness. Linear mixed effects models were used to examine differences in bees among regions, taking the form: *Response variable ~ region+Date+Trap+(1|Site:Replicate)*. The response variables examined were bee taxa richness and Shannon diversity. Tukey pairwise comparisons were performed using the *emmeans 1.7.4-1* package for all models to compare between regions.

## Results

Three sampling periods at our nine sites yielded 252 samples: 72 from the north, 107 from the central, and 73 from the south. From these samples, we identified 22,459 specimens: 9,304 from the north sites, 11,107 from central, and 2,048 from the south. In the central and southern regions, Diptera was the most abundant order. In the northern region, Hymenoptera was the most abundant order, due to a single emergence event where 6,032 chalcid wasps were collected, followed by Diptera (Table 1).

Traps at the southern sites produced the fewest unique taxa, followed by northern sites, although the two regions did not differ statistically from each other in richness. The central sites captured the most unique taxa, although this did not differ significantly from the richness captured at northern sites. There were no differences detected for Shannon diversity between any regions (Figure 4).

**Figure 4:**
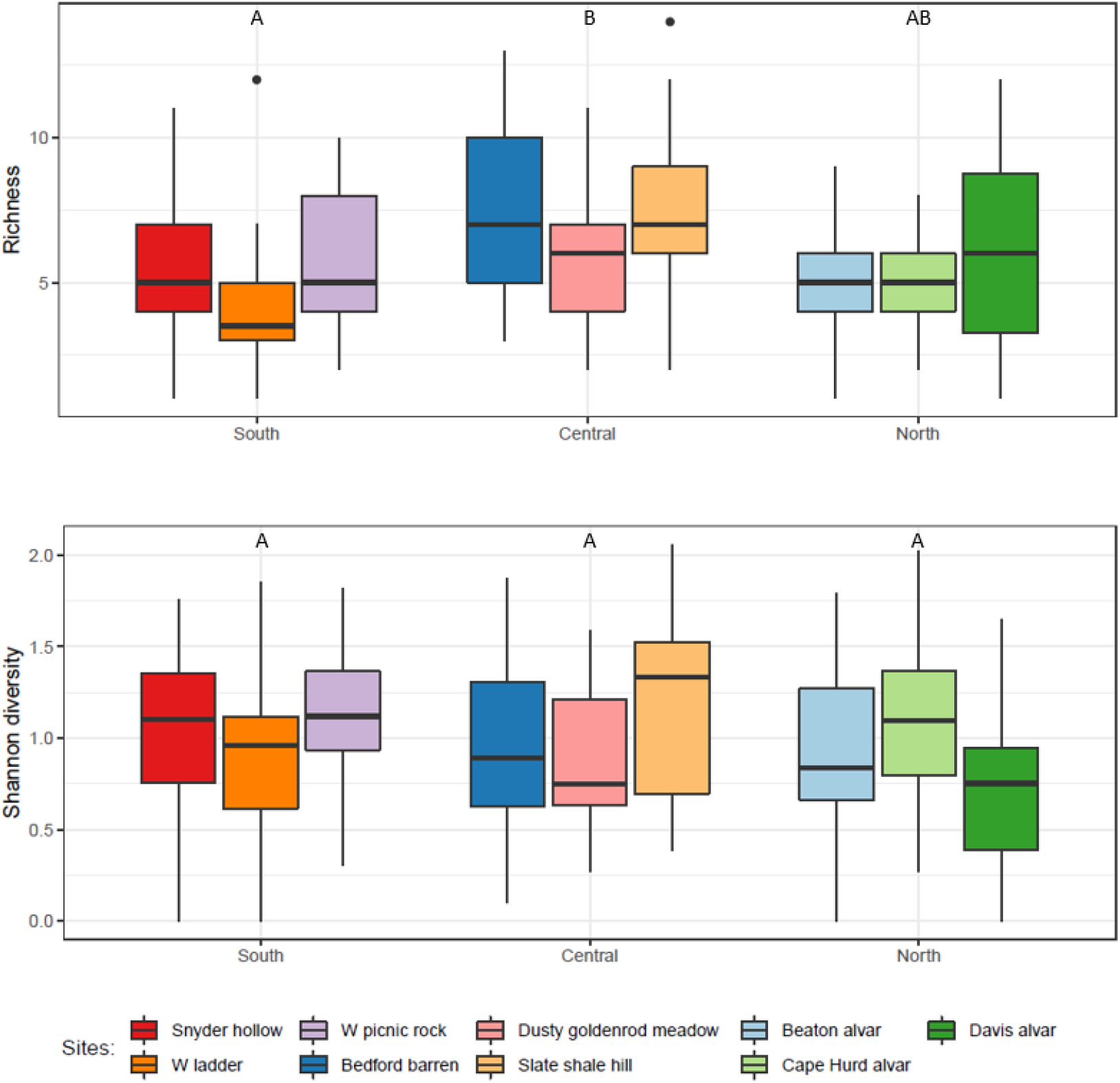
Box plots displaying taxa richness and Shannon diversity of insects, comparing sites by region. Letters shared indicate no statistical difference in estimated marginal means by Tukey method, P <0.05.

Non-metric multidimensional scaling (stress = 0.22) suggested strong overlap in insect communities between northern, central, and southern regions at this taxonomic resolution (Figure 5). When examined by sites within each region, the communities at all sites overlapped one another (Figure S1). The PERMANOVA testing for differences between regions was statistically significant (p = 0.001). Pairwise PERMANOVA found a statistically significant difference between southern and central regions (p = 0.01).

**Figure 5:**
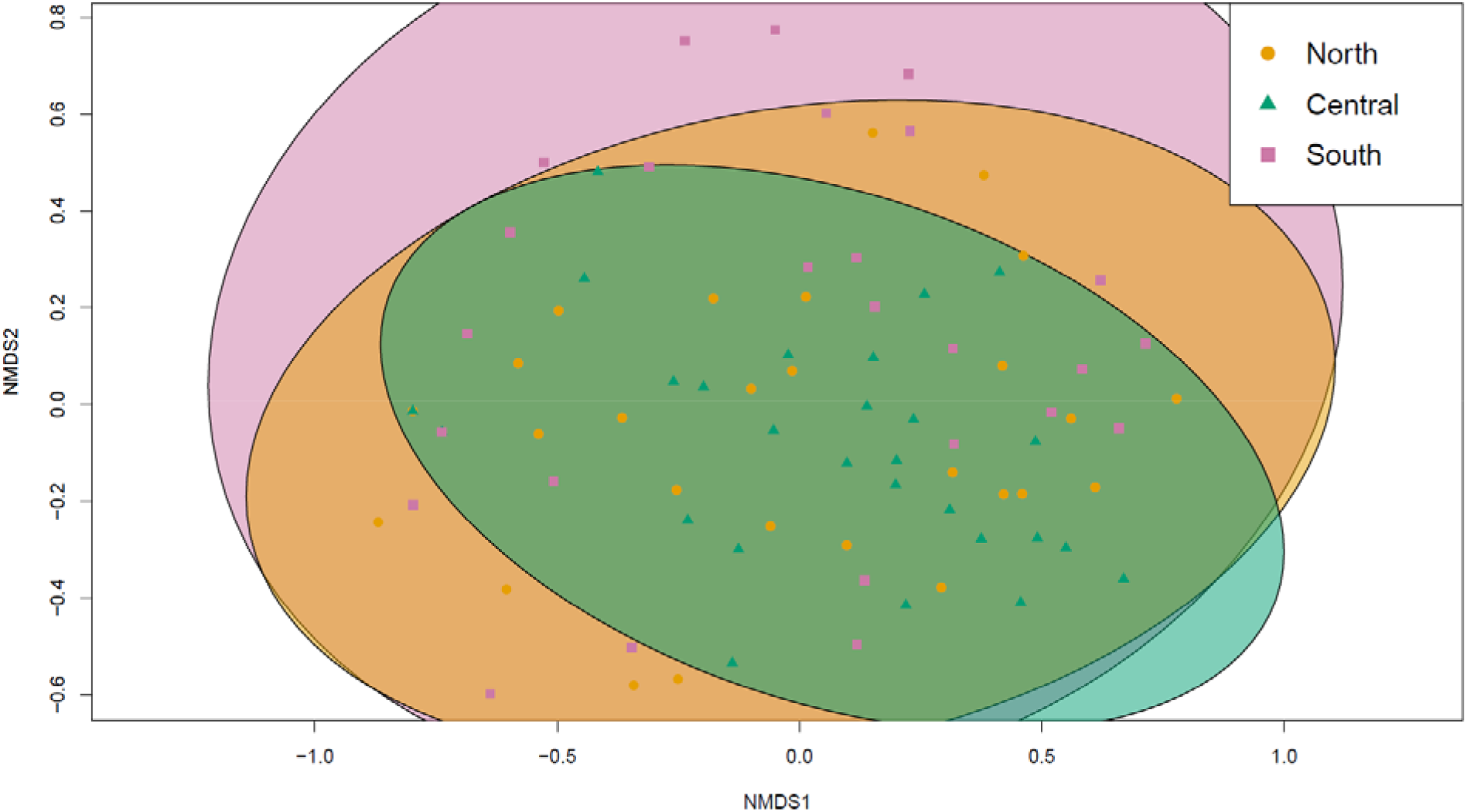
Non-metric multidimensional scaling figure (stress = 0.22) representing insect communities by region: North, Central, and South.

### Bees

Overall, there were 169 bees collected across all sites during our sampling period. We identified 151 caught in the bee bowls and ramp traps, excluding 18 individuals caught by the sticky cards as the glue damaged the specimens and obscured some identifying features. In the bee bowls and ramp traps there were 26 taxa. Of the identified specimens, 14 individuals from 5 taxa were collected in the northern region, 88 from 19 taxa in central, and 49 from 16 taxa in the southern. When compared with first order jackknife richness estimates, capture efficiency in the northern region was 74%, 71% in the central region, and 63% in the southern region (Figure 6). Region was not found to contribute significantly to bee community variation as there was no difference in bee richness or Shannon diversity between regions (Figure S2).

**Figure 6:**
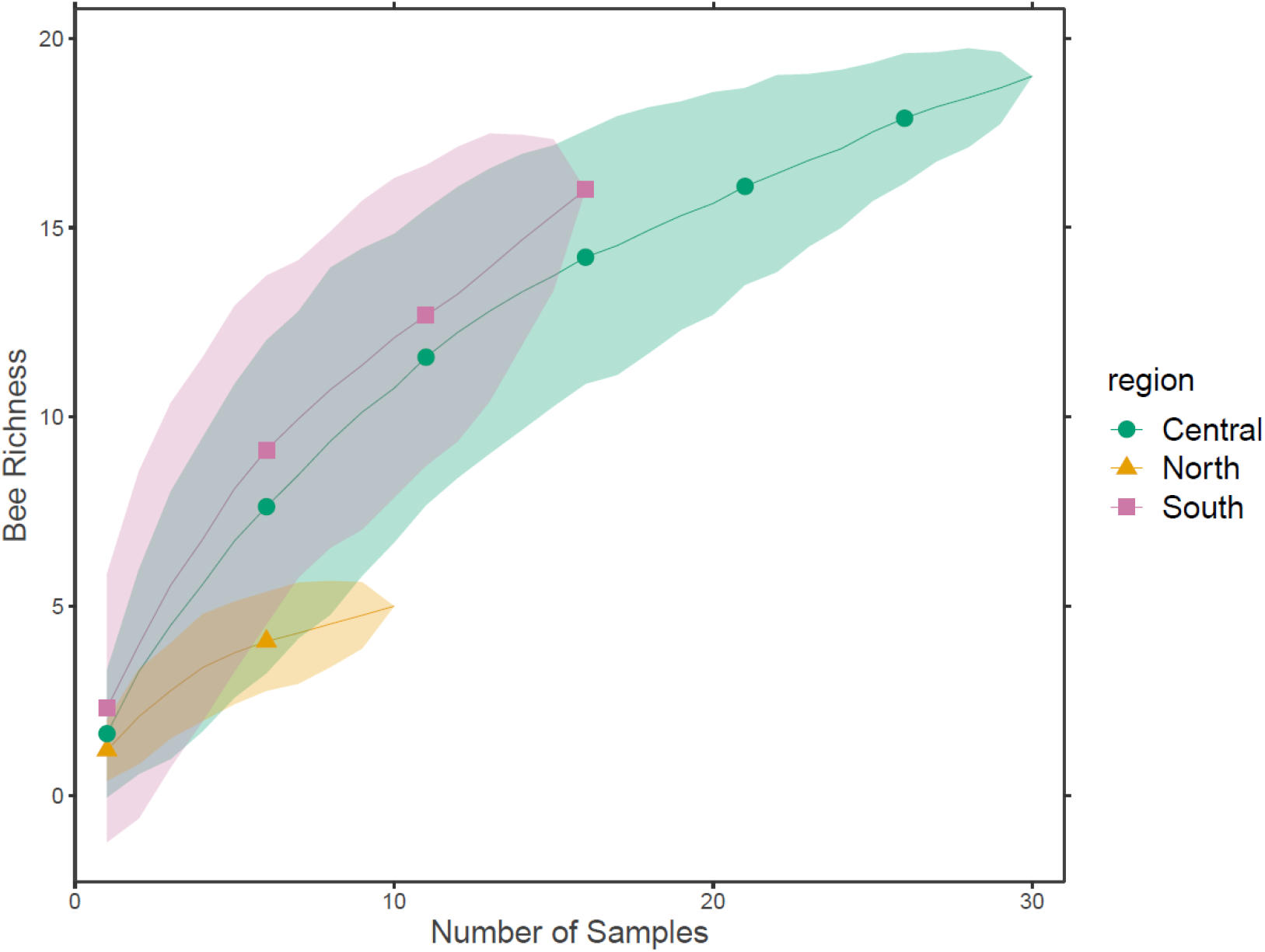
Accumulation curves for bee richness in each region. Shading represents standard deviation from the mean.

The single most abundant species in the north was *Augochlorella aurata* (Smith), but the genus *Lasioglossum* was more abundant overall. In the central region, *A. aurata* was the most abundant species. In the southern region, *Lasioglossum versatum* (Robertson) was the most abundant species, and there were 42 specimens in the genus *Lasioglossum* in total. One genus was unique to the northern region: Megachilidae: *Heriades*. Eight taxa were unique to the central region: Andrenidae: *Andrena*; Apidae: *Holcopasites*; Halictidae: *Agapostemon virescens* Fabricius, *Halictus ligatus* (Say), *Halictus confusus* Smith, *Lasioglossum coeruleum* (Robertson), *Lasioglossum tegulare* (Robertson), and *Lasioglossum zonulum* (Smith). Seven taxa were unique to the southern region: Apidae: *Apis mellifera* L., *Ceratina calcarata* Robertson; Halictidae: *Lasioglossum bruneri* (Crawford), *Lasioglossum cressonii* (Robertson), *Lasioglossum quebecense* (Crawford), *Sphecodes*; Megachilidae: *Megachile*.

### Plants

Regions varied substantially in plant species present. Overall, 53 vascular plant taxa were identified across all our sites. There were 21 taxa identified in the northern region, 24 in the central, and eight in the south. The only species recorded in more than one region was *Acer rubrum* L. (Red Maple), found in the central and southern regions (Table S2). Red maple was found at all three sites in the central region and two sites in the southern region.

## Discussion

This study aimed to compare insect biodiversity characteristics and trends among thin-soil environments across a latitudinal gradient in the Southeastern Great Lakes Region of North America. Our findings suggest that there is strong similarity among the insect communities within these habitats, at least when using typical sampling and community classification metrics. We saw no differences in Shannon diversity between any regions when compared over a three month span. However, the central region had greater insect taxa richness than the southern region. Visualization with NMDS showed complete overlap of insect community in all three regions, indicating similar community composition. The significant difference detected between the central and southern regions with the pairwise PERMANOVA suggests these differences were likely driven by the greater variability in community composition among sites in the southern region, while sites in the central region were more similar to each other. Similar trends were also observed when data were examined at the site level.

Bee analyses followed similar trends to the overall insect community analysis, as we observed similar taxa richness and Shannon diversity among regions. However, there were unique taxa collected within each region: one in the north, eight in the central, and seven in the south. It is important to note that the traps in the northern region captured a very low number of bees (14), compared to the central (87) and southern (49) regions, and between all the regions specimen numbers were relatively low, which should be considered when interpreting these results. However, there is strength in the higher resolution of identification of the bee specimens. With these data, we can identify some apparent associations of bee taxa with regions and sites. For example, *Augochlorella aurata*, a species in the family Halictidae, amounted to about 45% of specimens collected in the central region. All but two of these individuals were captured at one site, a sparsely vegetated hill with coarse slate and shale soil. *Augochlorella aurata* is a primitively eusocial, common, generalist, ground-nesting species (Wilson and Messinger Carril 2016). This species was caught in the northern and southern regions as well, but in much lower numbers. Two genera of cuckoo bee, brood parasites or kleptoparasites (Wilson and Messinger Carril 2016), were captured during the study. Cuckoo bees lay their eggs in the nests of other bee species, allowing their offspring to consume the resources meant for the host’s offspring, whom they typically kill (Wilson and Messinger Carril 2016, Litman 2019). Apidae: Nomadinae: *Holcopasites*, which typically parasitizes bees in the subfamily Panurginae, was unique to the central region. Halictidae: *Sphecodes*, which usually parasitizes other bees in their family, was unique to the southern region. The genus *Lasioglossum*, in the family Halictidae, is the largest genera of bees and notoriously difficult to identify (Gibbs 2018), however, 80% of *Lasioglossum* specimens were identified to species in our study. Half of the bees captured in the northern region were *Lasioglossum*, with three of the seven species collected belonging to the subgenera *Dialictus. Lasioglossum* dominated the southern region, amounting to over 85% of bees captured, most of these (43%) were *Lasioglossum (Dialictus) versatum*. *Lasioglossum versatum* is a primitively eusocial, generalist, ground-nesting species (Michener 1966). This species was captured at all three southern sites, on sandstone cliff bluffs. However, it was not unique to this region as specimens were captured in the central region as well, mostly at the meadow site.

During the three months of our study, we captured and identified 151 bee specimens belonging to 26 taxa overall. The northern region had the lowest bee richness and raw abundance and the central region had the highest bee richness and raw abundance. Though the bee bowls were the only trap targeting bees, we used the specimens collected in the ramp traps, targeting ground-dwelling insects, as well. A bee monitoring study that took place on non-thin soil environments in the county adjacent to our southern region sites (Washington County, OH, USA) captured 2,753 specimens containing over 130 species using bee bowls and Malaise traps in April-Oct 2013. When comparing the sampled species richness in only the bee bowls to jackknife 1 estimates, the capture efficiency was 73% overall, ranging from 70-74% at their sampling sites (Spring et al. 2017). On the Niagara peninsula, Ontario, Canada, between our northern and central regions, 15,733 bee specimens containing 124 taxa were sampled in April-Oct 2003 using pan traps (bee bowls), sweep and targeting netting. All three methods resulted in an estimated 84% capture efficacy, using ACE estimates. Pan traps alone captured 96 species, resulting in an estimated 75% capture efficiency (Richards et al. 2011). Although we collected a relatively low abundance of bees compared to these more intensive studies, using jackknife 1 richness estimates, our capture efficiency was comparable at 69% in total, ranging from 63-74% for the regions.

Overall, we captured few large-bodied bees, and notably no bumble bees, during the study, but that is most likely a result of the bias of bee bowls to collect smaller bodied genera (Wilson et al. 2008, Berglund and Milberg 2019, Portman et al. 2020). In future studies, adding hand-netting as a complementary sampling method could produce increased richness across sites and increase the comparability of bee taxa among these regions (Grundel et al. 2011, Prendergast et al. 2020). For example, Grundel et al. (2011) sampled bee communities in a variety of habitats in Indiana Nature Preserves using bee bowls and hand-netting techniques. They reported high abundances of larger species such as *Bombus impatiens* Cresson (Common Eastern Bumble Bee) and *Apis mellifera* L. (European honey bee) captured with hand nets and smaller-bodied families such as Halicitidae (sweat bees) captured in the bee bowls. We only captured one *Apis mellifera* during the study, but this is a common introduced, managed species found over most of the world (Butz Huryn 1997), and in abundance in other studies in the Great Lakes region (Spring et al. 2017, Rowe et al. 2020, Turo et al. 2021). Future investigators should consider identifying other biologically important taxa, such as beetles, to a higher resolution as other groups of insects may oy may not follow similar patterns as bees if identified to species.

The Davis alvar performed slightly differently from the other northern sites, as we observed lower Shannon diversity (Figure 3) and higher raw insect abundance. These differences may be attributable to large numbers of a parasitoid wasp (Trichogrammatidae) which emerged during our 12-14 August sampling period. Trichogrammatidae (superfamily Chalcidoidea) are known to parasitize eggs from several insect orders. Some genera, especially Trichogramma, are used for biological control of pest insects (Jalali et al. 2016). We captured just over 6,000 individuals that all appeared to belong to the same Trichogrammatidae morphological group, at only this site. When this emergence is removed from the data, the total number of Chalcidoidea collected in the northern region is much closer to that of the southern region (178 and 179, respectively). However, our analyses detected nearly identical patterns in comparisons of taxa richness and Shannon diversity with or without emergence data included. Without the emergence, the Davis alvar still had the greatest raw abundance of insects of the northern sites. Presence-absence data was used in the NMDS analysis so this emergence event would not influence the analysis of insect community composition.

From our observations, plant communities differed between regions with all three regions producing nearly unique species lists. There were no plant taxa present in all three regions. The only species we saw in more than one region was red maple, which was present in the central and southern regions. *Clinopodium arkansanum* (Nutt.) House (Limestone Calamint) was found at all three sites of the northern region, but absent from the central and southern regions. This species is associated with lakeshores and has been commonly found in flat rocky alvars near Lakes Huron and Michigan (Voss and Reznicek 2012, Cohen et al 2015). *Gaultheria procumbens* (American Wintergreen) and *Pinus virginiana* (Virginia Pine) were present at all three southern sites but absent from the central and northern regions. American wintergreen prefers sandy, acidic soils (Homoya and Namestnik 2022), which the sandstone soils of the southern region provide. Virginia pines are often found in soil derived from sandstone and their range overlaps with only the southern region (Snow 1965). The southern region, with its xeric, moss and lichen covered sandstone soils supported the lowest vascular plant richness.

Though the plant communities were unique in each region, we found overall similar insect community structure, perhaps due to their shared structural characteristic of thin soils. In general, insect communities were more similar in the north and central regions, with the southern region having lower insect taxa richness than the central region. Insect richness is often correlated with plant richness (Haddad et al. 2001), and we found the lowest vascular plant richness in the southern region as well. In this study, most insects other than bees were identified to order or family level which could have obscured some community patterns, and this caveat should be kept in mind when interpreting the findings.

The northern and central sites represent very different recent land-use histories but have the common history of both being within the extent of glaciation. The former two sites are situated well within the Great Lakes basin and the latter site is situated at the edge of this region, as it transitions into the Allegheny plateau. The history and severity of human disturbance at the sites also have the potential to drive insect community characteristics. One difference between the regions is their vehicular and pedestrian access and proximity to urban areas. The sites of the central region are proximate to urban areas while northern or southern region sites are rural and are on privately held nature preserve property. The central sites are open to the public, however not in areas typically with high foot traffic. Human disturbance has been well-established to affect insect abundance and diversity (Winfree et al. 2009, Quintero et al. 2010, Owens et al. 2020, Fenoglio et al. 2021, Raven and Wagner 2021, Wagner et al. 2021), with cities providing refuge to some insects, including pollinators (Hall et al. 2017). Though the central sites were located within preserved areas, those sites are still within the urban matrix of the greater Cleveland, Ohio area and have a history of human disturbance. Hemiptera was the second most abundant order in the central region due to the large number of aphids, a common crop pest, captured (Miller and Foottit 2009, Emden and Harrington 2017). This and the higher overall numbers of insects collected in the central region could be a result of previous disturbance.

Thin-soil environments, while considered important rare habitats, may also serve a more utilitarian role as model systems. These habitats have been studied as biotic and abiotic templates for urban green, or vegetated, roof design because of their structural similarities (Lundholm 2006, Lundholm and Walker 2018). Extensive green roofs are a type of living architecture in which plants are intentionally grown on top of a human-built structure in shallow (typically 15 cm or less) growing medium (Getter and Rowe 2006). Like the natural thin-soil environments, they may face similar weather patterns as they are in an exposed area: varying precipitation conditions, intense solar radiation, and wind. Thus, our expectations for biotic function in these environments should too be modeled after natural systems with similar traits. Green roofs have the potential to provide urban areas with many services, including stormwater retention, reduced building energy consumption, habitat for organisms, and more depending on how it is designed (Dunnett and Kingsbury 2004). Plants and other physical structures from natural thin-soil environments may be used to design green roofs that feature native plants and landscapes similar to those found outside of the urban area (Lundholm 2006, Lundholm and Walker 2018). These habitat patches may aid in conservation of plants and insects found in those natural ground-level environments and increase certain ecosystem services, such as pollination and pest control, for the city (Kadas 2006, Colla et al. 2009, MacIvor and Lundholm 2011).

## Conclusions

Thin-soil environments, characterized by thin or absent soils atop hard substrates, are unique and globally rare environments (Reschke et al. 1999, Cohen et al. 2015, Neufeld et al. 2018). Due to the nature of their soils, these open habitats can be quite harsh from an abiotic standpoint and thus home to unique communities of plants and animals (Albert et al. 1995, Comer et al. 1997, Reschke et al. 1999, Neufeld et al. 2018, McMullin 2019). While previous work has examined the composition of these habitats (Albert et al. 1994, 1995, Comer et al. 1997, Bouchard et al. 1998, 2005, Reschke et al. 1999, Albert 2006, Cohen et al. 2015, Neufeld et al. 2018, McMullin 2019), in this study, we examined how these environments shape similarities in insect communities. This study leverages a unique design that encompasses a large area of the Southeastern Great Lakes Region, spanning over 600 km north to south.

By examining the plant and insect communities in three separate regions of this area of North America, we found that overall insect community composition and biodiversity characteristics were similar. Though some variation in taxa richness was detected, Shannon diversity was similar across all three regions. The southern region had the lowest insect taxa richness and the lowest vascular plant richness. All regions contained unique plant taxa, with only one species being found in more than one region. Bee identity was similar across all regions, finding no differences in biodiversity metrics and similar community composition, though each region contained unique taxa.

This study provides taxonomic information about the insect, particularly bee, and plant communities in thin-soil environments in this region. This knowledge supports conservation and management efforts of rare thin-soil habitats such as alvars. Furthermore, this study provides insights into how communities of insects assemble in habitats with these structural characteristics. If these habitats are modeled by architects or urban planners for green roofs or other urban infrastructure, insights can be used to create realistic expectations for biodiversity outcomes in the built environment.

## Data and code

https://github.com/katiemmanning/Thin-soil

## Notes

https://github.com/katiemmanning/Thin-soil

### Competing Interest Statement

The authors have declared no competing interest.

### Summary of Updates

This manuscript contains additional figures and tables, revised statistical methods, and additional discussion of the results.

