## Supplemental materials for "Characterizing insect communities within thin-soil environments"

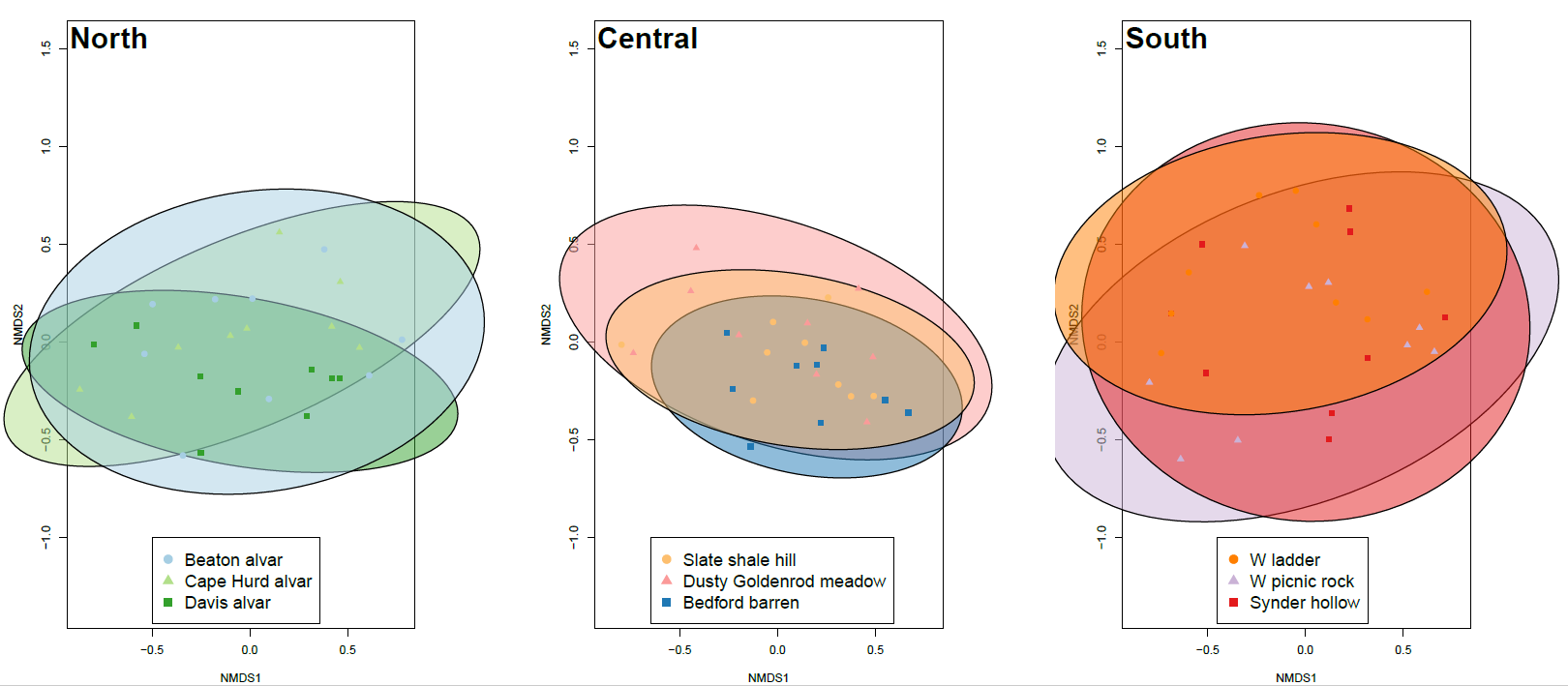

**Figure S1: Non-metric multidimensional scaling figure (stress = 0.22) representing insect communities by site within the regions.**

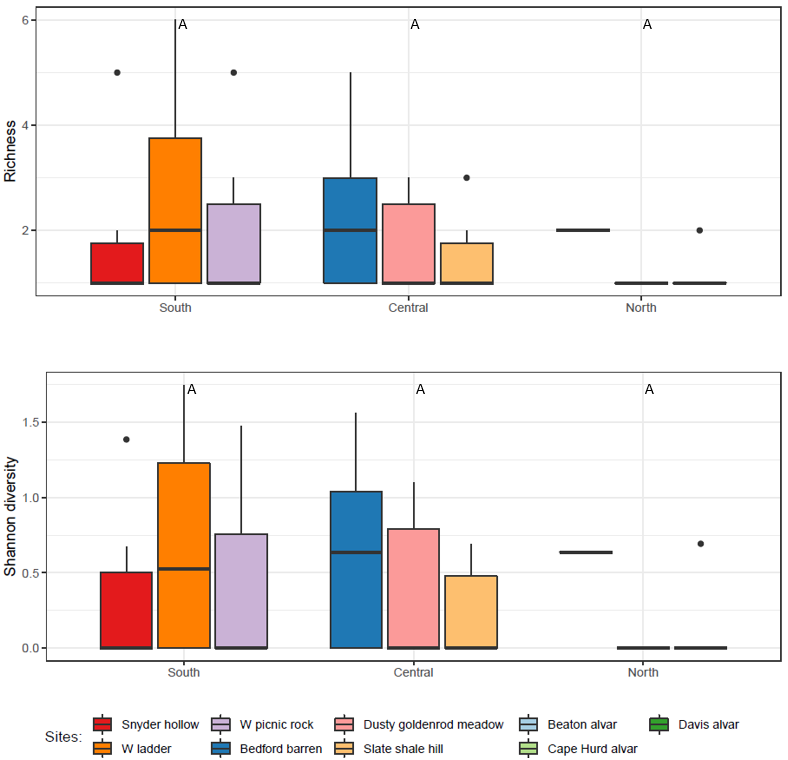

**Figure S2: Box plots displaying insect taxa richness and Shannon diversity of bees, comparing sites by region. Letters shared indicate no statistical difference in estimated marginal means by Tukey method, P <0.05.**

**Table S1: Overview table of insect sampling**

|  |  |  | **Number of traps** | | |
| --- | --- | --- | --- | --- | --- |
| **Region** | **Site** | **Classification** | **Sticky cards** | **Ramps** | **Bowls** |
| North | Davis alvar | Alvar | 4 | 3 | 3 |
|  | Beaton alvar | Alvar | 3 | 2 | 2 |
|  | Cape Hurd alvar | Alvar | 3 | 2 | 2 |
| Central | Dusty goldenrod meadow | Meadow | 6 | 3 | 3 |
|  | Bedford barren | Barren | 6 | 3 | 3 |
|  | Slate shale hill | Barren | 6 | 3 | 3 |
| South | "The W" - ladder | Barren | 4 | 2 | 2 |
|  | "The W" - picnic rock | Barren | 4 | 2 | 2 |
|  | Snyder hollow | Barren | 4 | 2 | 2 |

**Table S2: Vascular plant taxa identified in each region**

| North | Central | South |
| --- | --- | --- |
| *Anticlea elegans* | *Acer rubrum* | *Acer rubrum* |
| *Arctostaphylos uva-ursi* | *Achillea millefolium* | *Amelanchier aborea* |
| *Asplenium trichomanes* | *Amelanchier* sp. | *Carex* sp. |
| *Carex eburnea* | *Apocynum* sp. | *Gaultheria procumbens* |
| *Carex richardsonii* | *Aster* spp. | *Gaylussacia baccata* |
| *Clinopodium arkansanum* | *Danthonia spicata* | *Photinia melanocarpa* |
| *Coreopsis lanceolata* | *Daucus carota* | *Pinus virginiana* |
| *Dasiphora fruticosa* | *Elymus* sp. | *Vaccinium pallidum* |
| *Deschampsia cespitosa* | *Hieracium* sp. |  |
| *Dichanthelium implicatum* | *Juncus effusus* |  |
| *Houstonia* sp. | *Nyssa sylvatica* |  |
| *Hypericum perforatum* | *Pinus strobus* |  |
| *Hypericum prolificum* | *Polygala nuttallii* |  |
| *Juniperus horizontalis* | *Potentilla* sp. |  |
| *Larix laricina* | *Prunus* sp. |  |
| *Minuartia michauxii* | *Quercus rubra* |  |
| *Packera paupercula* | *Rhynchospora* sp. |  |
| *Picea marinana* | *Rubus* sp. |  |
| *Solidago hirsuta* | *Schizachyrium scoparium* |  |
| *Solidago ptarmicoides* | *Solidago* spp. |  |
| *Thuja occidentalis* | *Sphagnum* sp. |  |
|  | *Spiraea tomentosa* |  |
|  | *Tsuga* sp. |  |
|  | *Viola* spp. |  |
